## Supplementary Figures with Caption for "Neural manifolds for odor-driven innate and acquired appetitive preferences"

**Department of Biomedical Engineering, Washington University in St. Louis**

**Supplementary Figures**

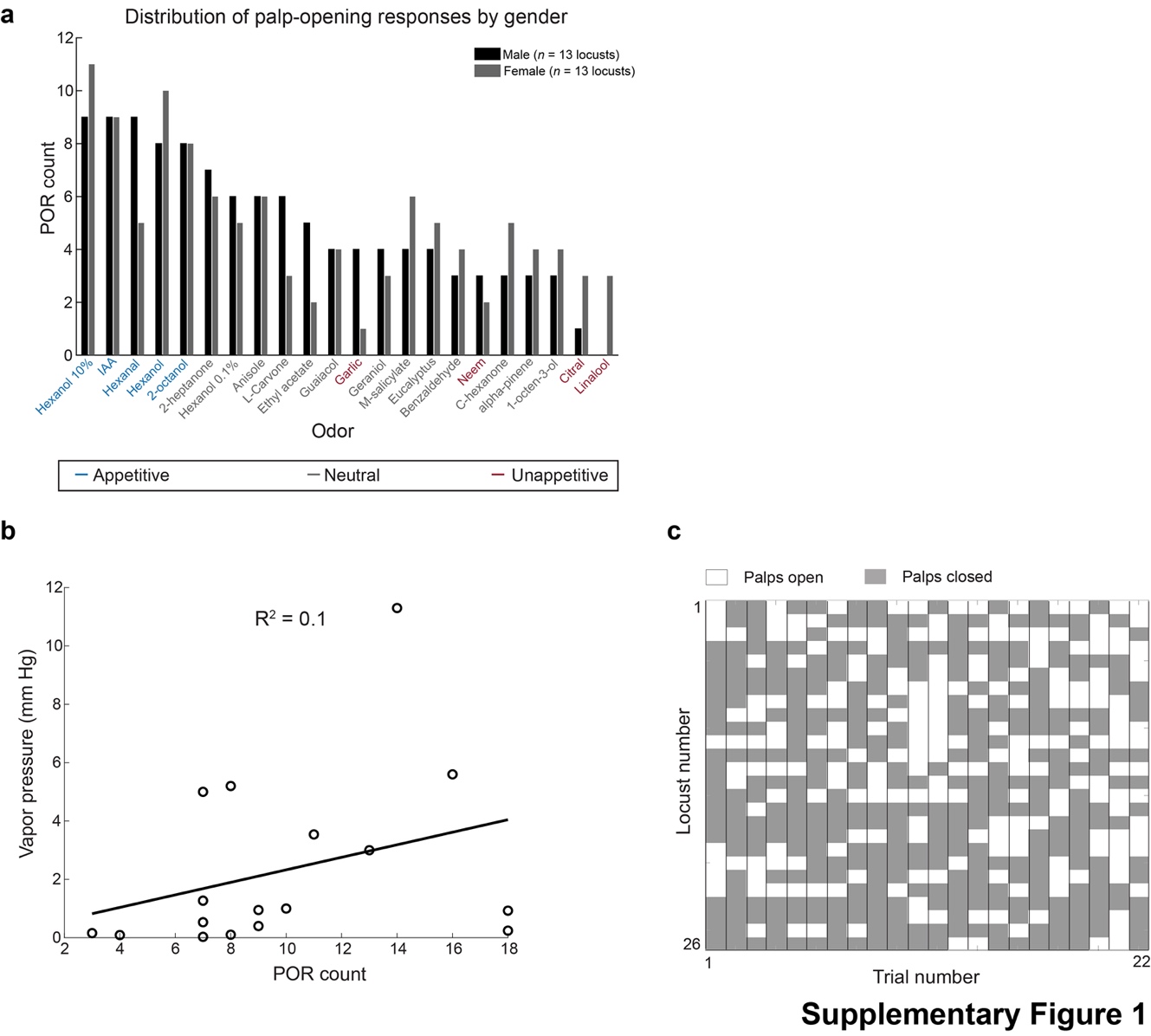

**Supplementary Figure 1**: Additional controls for assaying innate appetitive preference

**a)** Palp-opening responses (PORs) to the odor panel do not show any significant gender-based differences (t-test, *p*>0.1 for all pairwise comparisons). We recorded PORs from 13 male and 13 female locusts to measure innate valence. PORs were sorted from highest to lowest based on the male group responses (black bars), and corresponding female locust PORs (dark gray bars) are shown next to them. The odors are colored using the same convention used in **Fig. 1c**.

**b)** Similar plot as in **Fig. 1d** but without ethyl acetate. Ethyl acetate has a reported vapor pressure that is much higher than all other odors used, and hence we repeated the analysis in **Fig. 1d** without the outlier. The R^2^ for this analysis is 0.1, which still indicates a very poor correlation between vapor pressures and the observed palp-opening responses.

**c)** The PORs recorded for every locust are shown as a function of trial number. Similar convention as **Fig. 1b**, where each row corresponds to a single locust and there are twenty-two trials one for each odorant in the panel. White boxes indicate a palp-opening response, gray boxes indicate no PORs. A summary of this data is presented in **Fig. 1d**.

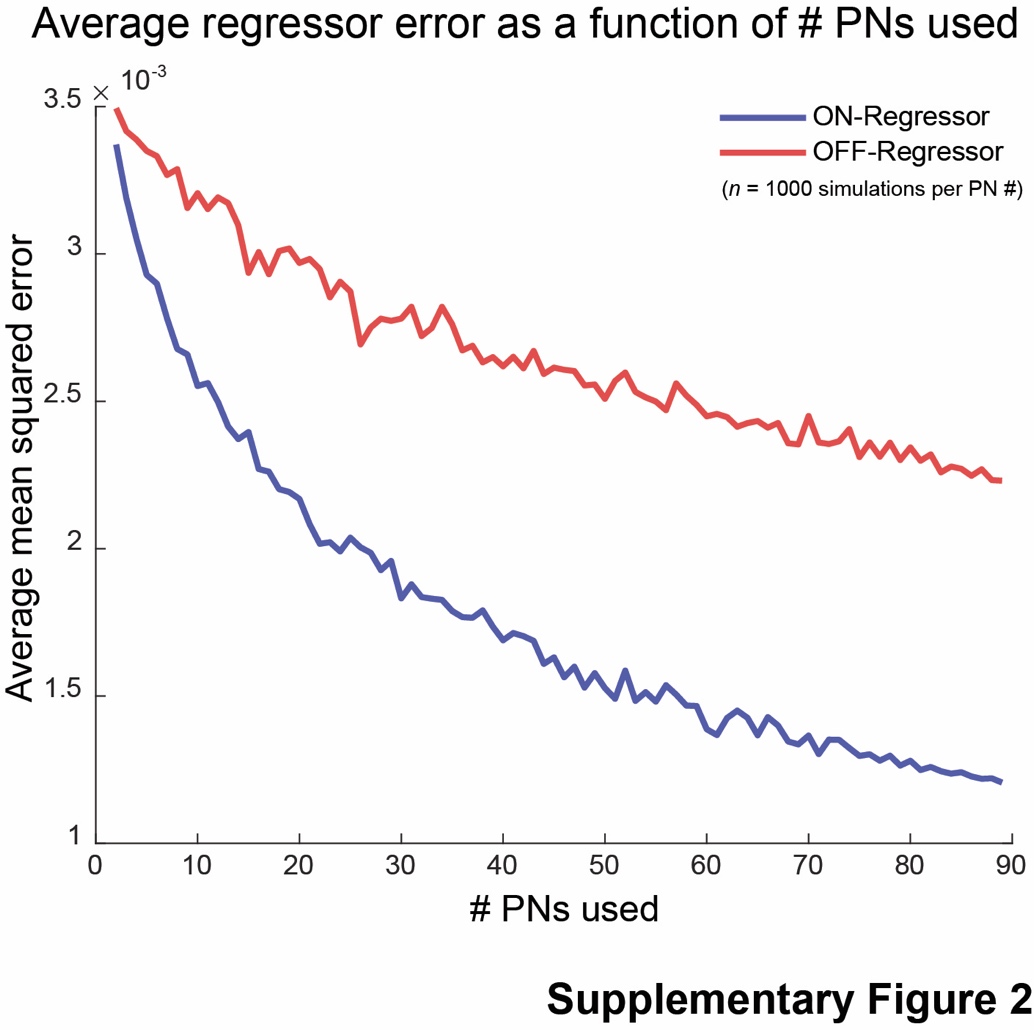

**Supplementary Figure 2**: Monte Carlo simulations to quantify the performance of the linear regression approach as a function of the number of projection neurons used to predict the behavioral preference indices. Note that the performance of both ON- (blue) and OFF- (red) regressors are plotted to allow comparison. For each *n*, we performed 1000 such simulations and obtained the average mean squared error for the predictions across all simulations. As can be seen, the average error goes down with an increase in the PN count and appears to saturate when around eighty PNs are used for the analyses.

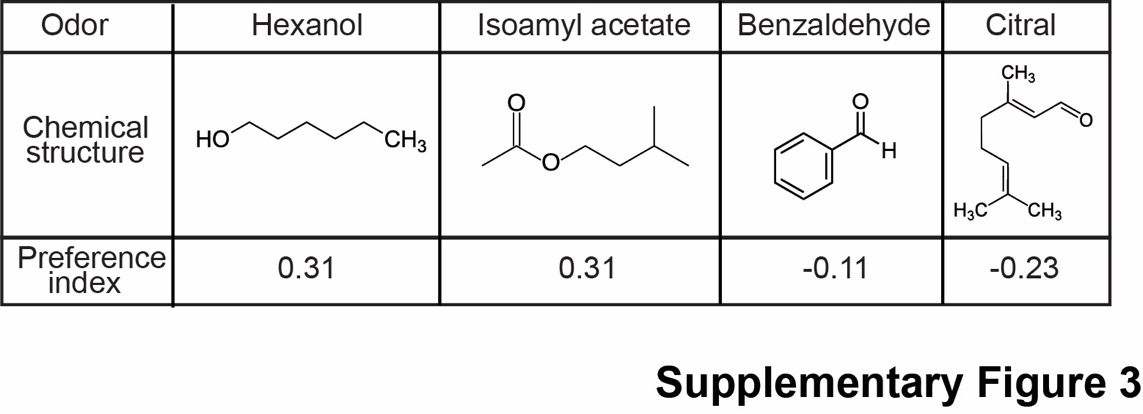

**Supplementary Figure 3**: Diverse odorants used for Pavlovian conditioning assays

We used 4 odors – hexanol, isoamyl acetate, benzaldehyde, and citral for all the appetitive conditioning assays. As can be seen here, these 4 odorants have very diverse chemical structures with unique functionalities, as well as diverse innate preference indices (from **Fig. 1**).

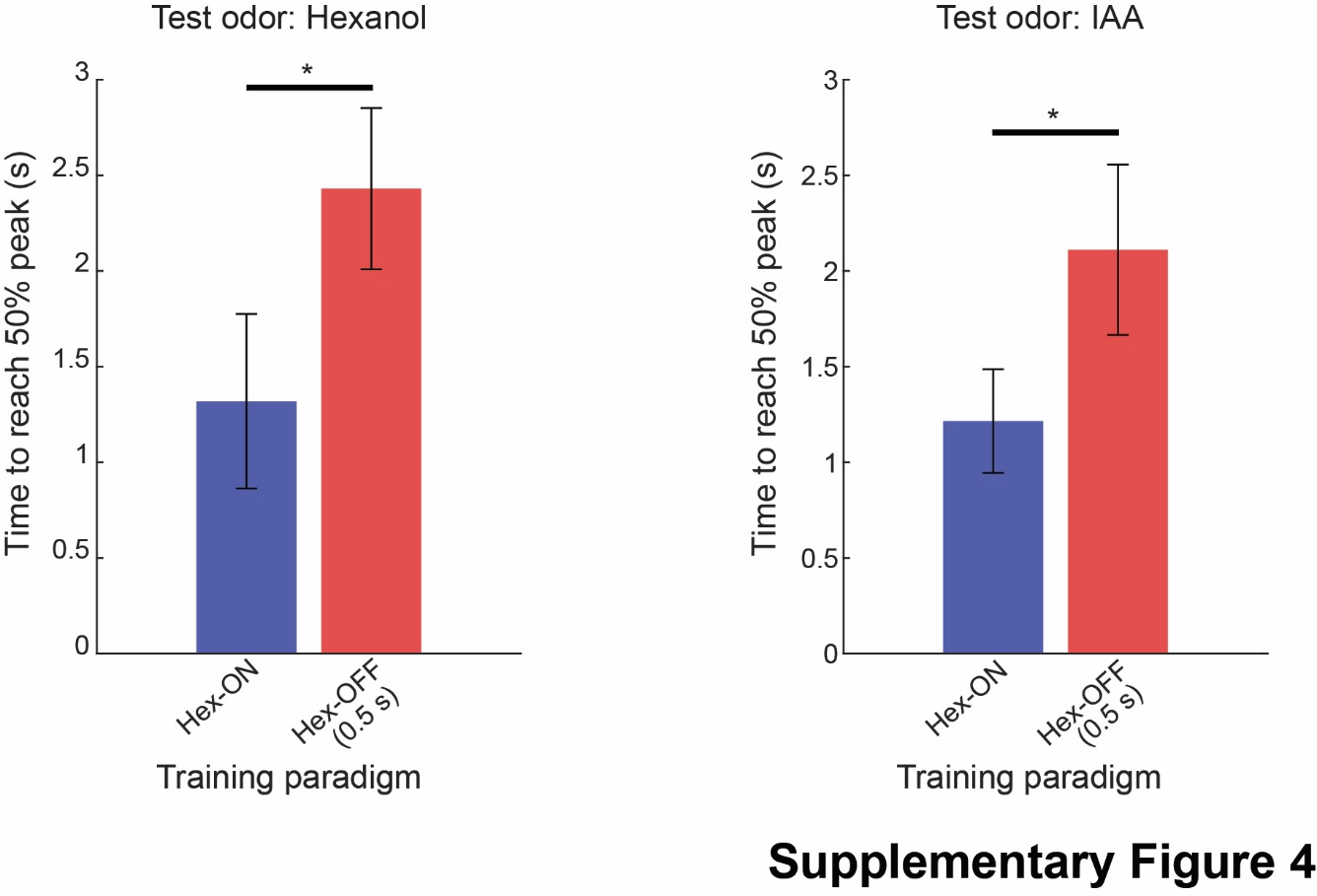

**Supplementary Figure 4**: ON- and OFF- conditioning produce temporally distinct responses

Latency of locust PORs to hexanol (left) and isoamyl acetate (right) are shown. Response latency here is defined as the time taken by locusts to reach 50% of the peak palp separation (time = 0 along the y-axis indicates odor onset). Each bar plot shows the mean latency across locusts, and error bars indicate s.e.m. For each odorant tested, POR latency for two groups of locusts either trained using hexanol ON-training paradigm (blue bars) or OFF- training paradigm (red bars) are shown for comparison. For this analysis, we used only those locusts that had significant responses to the test odorant (refer **Fig. 6b** for fractions; see **Methods**). For both odors, locusts used in the hexanol OFF-training paradigm were significantly slower in opening their palps (* indicates *p* < 0.05, one-sided t-test).

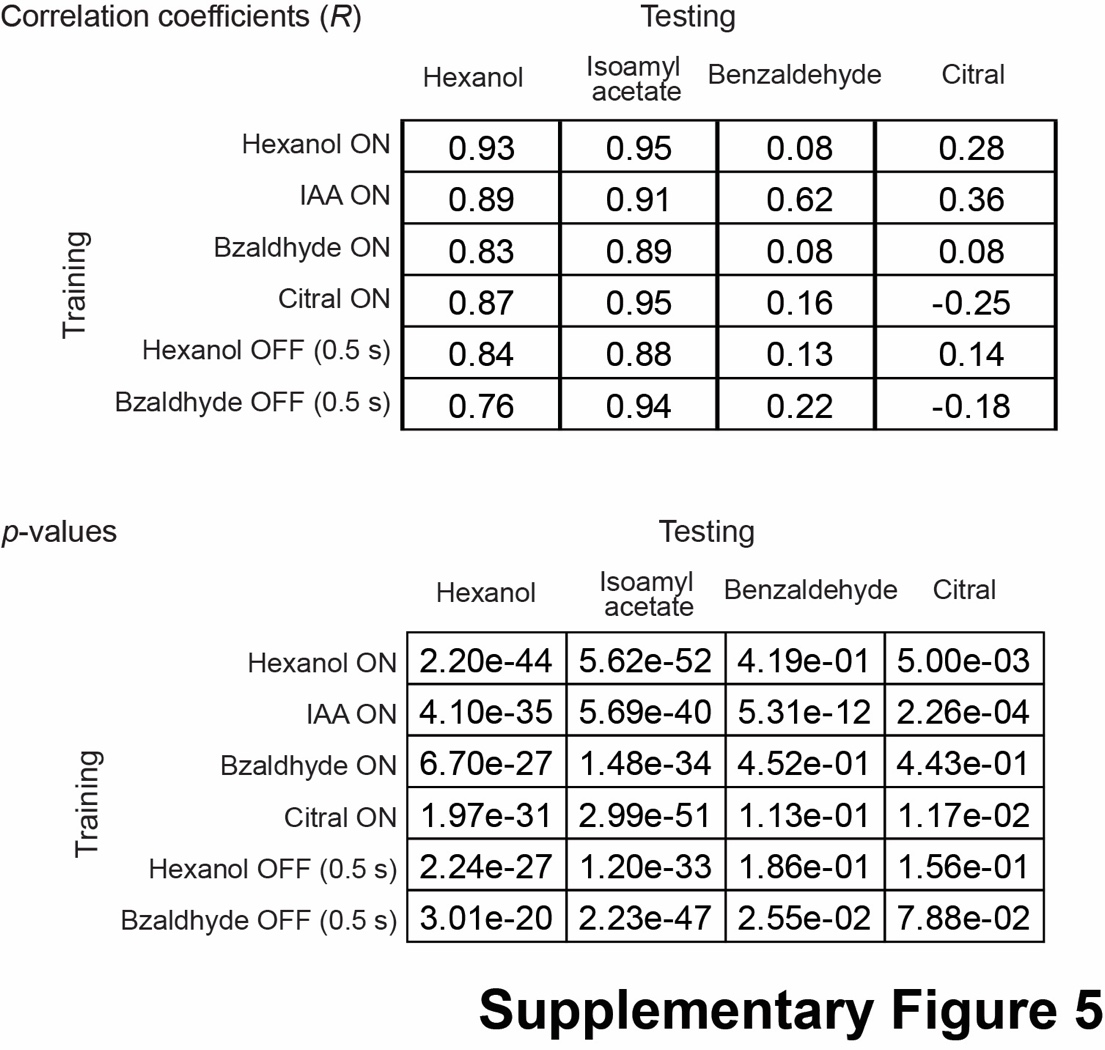

**Supplementary Figure 5**: Quantifying model performances to predict mean POR traces

The two tables show the correlation between the predicted POR versus the observed behavioral response dynamics (R, top table) and significance (*p*-value, bottom table) (red traces in **Fig. 6a**). Similar to the convention in **Fig. 6a**, each row corresponds to one training paradigm and each column shows one test odor.

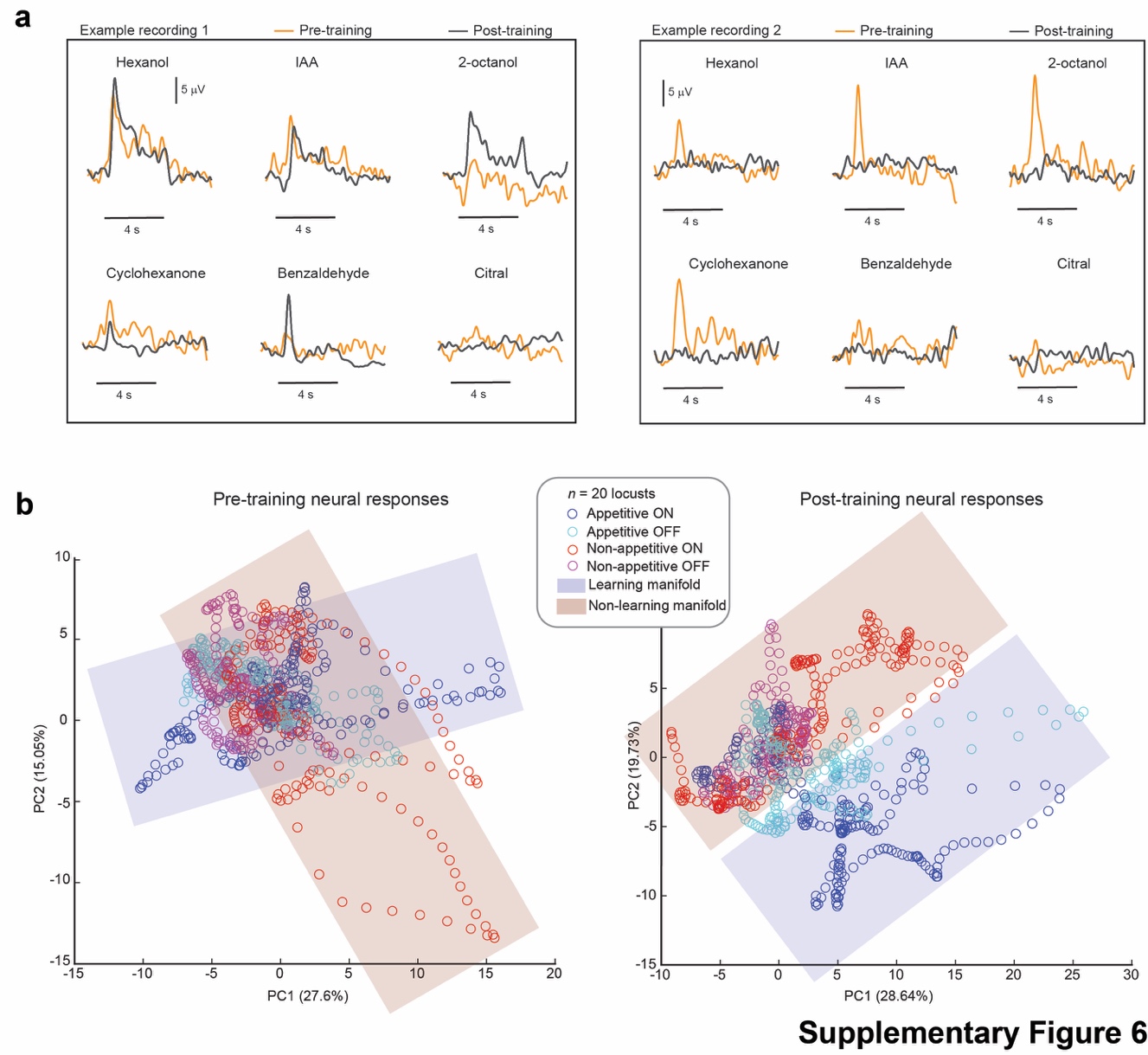

**Supplementary Figure 6**: Neural responses manifolds in behaving locusts

**a)** Representative recordings showing the total odor-evoked responses (see **Methods**) to the odor panel before (orange) and after training (gray). In example 1 (left panel), hexanol was used as the training odorant, and in example 2, benzaldehyde was used as the training odorant (right panel). In each panel, the top row contains the appetitive odorants and the bottom row contains stimuli that were non-appetitive. Black bars below the plots indicate 4 s of odor presentation.

**b)** PCA visualization showing ensemble neural responses during both the ON- and the OFF- periods for all 6 odors are shown along the top 2 principal components (*n* = 20 locusts; see **Methods**). The data points were colored as follows: blue – appetitive odorants ON responses, cyan – appetitive odorants OFF responses, red – non-appetitive odorants ON responses, and magenta – non-appetitive odorants OFF responses. Variances in odor-evoked responses of appetitive odorants were not uniformly distributed but confined to a subspace and are schematically shown as using a linear plane (plane colored in blue that encompasses appetitive ON and appetitive OFF neural ensembles). Similarly, non-appetitive odorants ensemble responses were confined to a distinct neural manifold schematically shown in red. Note that the two subspaces became less overlapping post-training (right panel versus left panel).

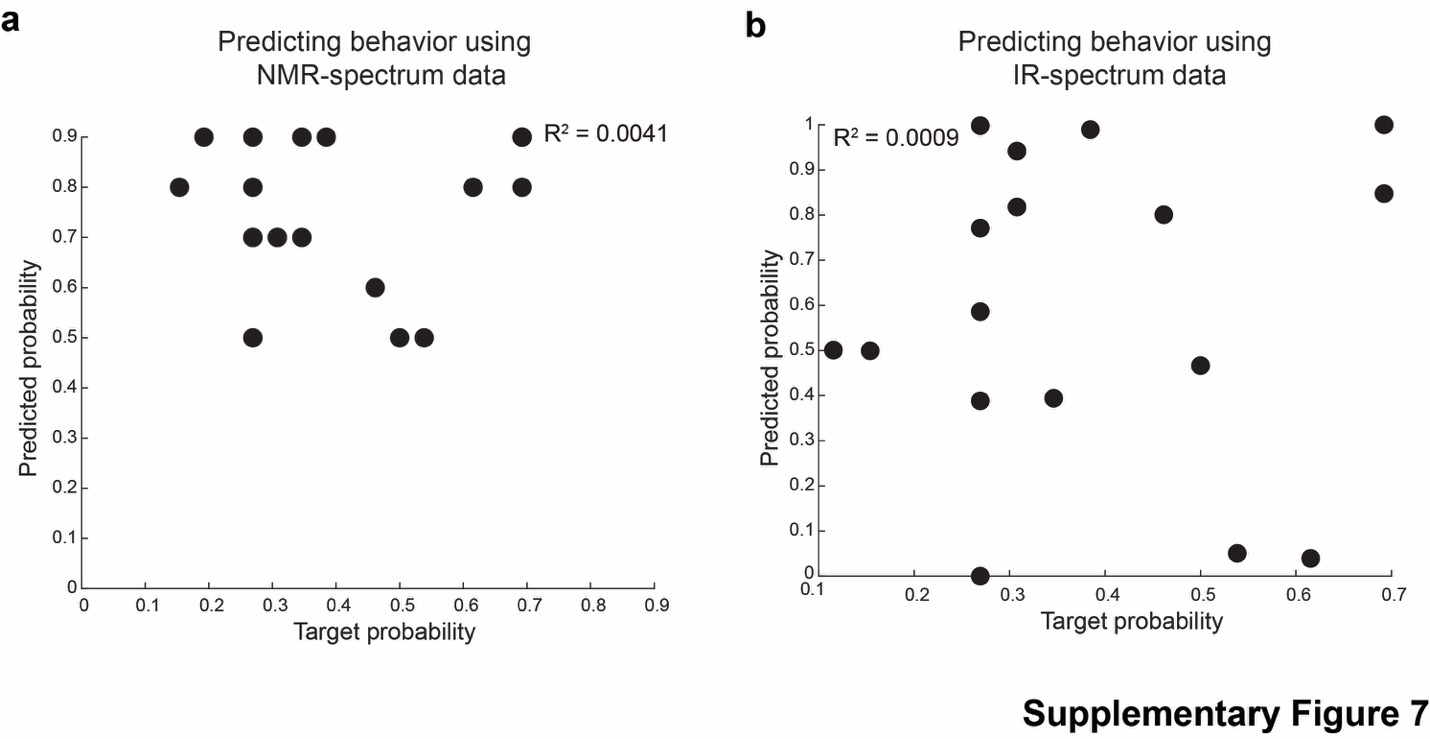

**Supplementary Figure 7**: Mapping chemical features/properties onto behavioral valence

**a)** We obtained NMR-spectrum data^53^ for 16 odors (all at 1% concentration) in our panel. Using an approach similar to that in **Fig. 4**, we trained linear regressors to predict the valence of odorants in the panel based on their NMR-spectra. The plot shows the actual POR probability along the x-axis and the predictions from the regressors along the y-axis. The predictions were poor and had a calculated R^2^ value of 0.0041.

**b)** Similar approach as in panel **a**, but using IR-spectrum data obtained for 17 odors^53^. The predictions were again poor and had a calculated R^2^ value of 0.0009.
